# Microbiome variations in osteoarthritis reflect aging and metabolic factors, not the disease

**DOI:** 10.1101/2025.06.24.661261

**Authors:** Kajetana Bevc, Lukas Malfertheiner, Samuel Neuenschwander, Van Du Tran, Marco Pagni, Kateryna Pantiukh, Madis Jagura, Teemu Niiranen, Rob Knight, Veikko Salomaa, Aki Havulinna, Elin Org, Kari K Eklund, Gonçalo Barreto, Marcy Zenobi Wong, Christian von Mering

## Abstract

The gut microbiome is crucial for human health. Its disruption has been linked to several chronic diseases, including joint disorders. The gut–joint axis has been implicated in the pathogenesis of osteoarthritis (OA), but conflicting findings and study limitations have led to uncertainty regarding the role of microbiota. We conducted a multi-cohort gut microbiota analysis in 1,395 OA patients from four European cohorts (Lifelines, EstMB, FINRISK 2002, TwinsUK), using stringent exclusion criteria and matched controls. When assessing microbial diversity, taxa, functional gene profiles, and gut permeability biomarkers, no significant differences were found between OA and controls. Although this does not exclude a causal contribution of the microbiota in the gut-joint-axis, its dysbiosis does not seem to be linked with OA disease progression. Instead, age and BMI appeared as the main contributing factors to microbiome changes. Microbiome studies in complex diseases often face challenges such as small sample sizes, batch effects, and limited ability to match appropriate controls, particularly in single-cohort designs. By combining data from multiple large cohorts, we were able to mitigate these limitations and provide a more robust assessment of microbiome association with OA. Our findings emphasize the need for rigorous study design in microbiome research and challenge the OA-gut dysbiosis hypothesis.

## 1. Introduction

The human body contains roughly the same number of bacterial cells as human cells, with the vast majority of bacteria residing in the colon^1^. The gut microbiome influences both health and disease as it is responsible for various metabolic and immunological functions^2,3^. The microbial species present in the gut as well as their diversity differ remarkably between individuals^4–6^ and are influenced by environmental and physiological factors such as diet, age and sex^6–9^. Therefore, there is no universal healthy microbiota fingerprint. However, in a state of dysbiosis, consistent signals driving pathology emerge; low bacterial diversity in the gut is linked to obesity and inflammatory bowel disease (IBD) as well as to a leaky gut^10,11^.

Gut permeability is the result of tight junction integrity disruption in the gut epithelium. This intermittent leakage of various bacteria and bacterial products into the circulation has been described as sufficient to trigger the onset and flares of diseases such as systemic lupus erythematosus and type 1 diabetes^11,12^. Gut leakiness is often assessed by serum zonulin, lipopolysaccharide (LPS), LPS binding protein (LBP), fatty acid binding protein2 (FABP2) levels and soluble CD14^13–15^. Zonulin is a reversible tight junction regulator recognized as a biomarker of intestinal permeability^16,17^. Whereas LPS is a class of bacterial molecules that is bound by LBP in human serum, both signifying higher levels of intestinal permeability^18,19^. Serum levels of Zonulin and LBP were reportedly increased in erosive hand osteoarthritis (OA) compared to non-erosive hand OA^20^. Karim et al also reported increased serum Zonulin in knee OA patients compared to controls and negatively correlated its concentration to functional performance, suggesting the role of zonulin in OA-induced physical disability^21^. From such studies it has been hypothesized that leaky gut is the underlying mechanism in the gut-joint axis^2^.

The gut-joint axis has been proposed to play a role in both OA and rheumatoid arthritis (RA), as LPS and bacterial DNA have been found in the synovial fluid, synovium and cartilage of these patients^12,22^. OA is a complex multi-tissue disease of the joints affecting 595 million people in 2020 with the numbers projected to increase for 74,9% by 2050 for knee OA^23^ due to the increasing prevalence of obesity and an aging population, both of which are major risk factors for OA. Another risk factor is sex, with older women with higher weight being disproportionately affected^23–25^. OA symptoms are debilitating and result in substantial decrease in the quality of life for the patients. Thus, OA poses a great socioeconomic burden costing an estimated US$80 billion in healthcare in the USA in 2016 alone^23^. Previously described as a “wear and tear” disease characterized by low grade inflammation, recent works point to the involvement of the gut microbiome^26,27^. Increased abundance of *Streptococcus* sp. and Firmicutes phylum were shown to be more abundant in OA patients’ microbiota^26^. The *Clostridium* genus, part of the Firmicutes phylum, particularly *Clostridium leptum* have been shown to be more abundant in OA individuals in several studies^28,29^. Interestingly, one study also reported a lower abundance of *Clostridium boltae* and suggests that this alters microbial bile acid formation and affects OA progression^30^. Firmicutes/Bacteroidetes ratio was also suggested to be higher in OA individuals^22,29,31^. However, other studies found a higher prevalence of the *Bacteroides* genus in OA patients ^22,28^. Beyond microbial diversity, there have been reports on increased serum levels of LBP and Zonulin in erosive hand OA^20^. Huang et al. found that increased serum levels of LPS and LPS-binding protein (LBP) were associated with knee osteophyte severity and abundance of activated macrophages in the synovium^19^. However, our study showed that systemic increase of LPS did not induce OA-like symptoms in rats and that LPS concentration in the synovial fluid did not differ significantly between OA and control^32^. Notably, Loeser et al also did not find any significant differences in microbiota composition between OA and control groups^33^. The poor clinical efficiency of doxycycline, an antibiotic with an effect on cartilage, in OA treatment also questions the involvement of bacteria in OA pathogenesis^34^.

Accumulating evidence has demonstrated that age, sex, and obesity, all known OA-risk factors, significantly contribute to gut dysbiosis, which poses the question if the purported association of gut dysbiosis to OA in previous population studies was an independent association^2,8,9,35^. Furthermore, the diverse environments in which humans live also likely strongly confound microbiome analyses^36^. This is further complicated by non-comparable sampling designs between cohorts and studies and by differing availability of metadata, such as use of different medications^8^. Additionally, many of the studies were performed with small patient subsets and did not account for different factors that are known to affect the gut microbiome, such as gut inflammation, pregnancy, and certain medications^37–39^. The heterogeneity of sampling and analysis methods, different taxonomic levels assessed, and the lack of healthy controls in some of these studies make it difficult to identify a causal link^29^.

Here, we elucidated the role of microbiota in OA and performed a well-controlled multi-cohort analysis of gut microbiota composition of patients with OA. OA samples were compared to matched healthy subjects with stringent exclusion criteria. This study focuses both on the characterization of microbiota phenotypes in OA as well as assessing the gut permeability.

## 2. Methods

### Cohorts

The cohorts we used were Lifelines, EstMB, FINRISK 2002 and TwinsUK.

Lifelines is a multi-disciplinary prospective population-based cohort study examining in a unique three-generation design the health and health-related behaviours of 167,729 persons living in the North of the Netherlands^40^. It employs a broad range of investigative procedures in assessing the biomedical, socio-demographic, behavioural, physical and psychological factors which contribute to the health and disease of the general population, with a special focus on multi-morbidity and complex genetics.

The Estonian Microbiome Cohort (EstMB) is a volunteer-based population cohort established in 2017, that currently includes over 2,500 adults across Estonia. A detailed overview of the EstMB, including omics and phenotypic data availability, is described in Aasmets & Krigul et al^41^. In brief, EstMB cohort includes 1,764 women (70.3%) and 745 men (29.7 %), aged between 23 and 89. Extensive information was collected on the EstMB participants, including electronic health records and the participants’ anthropometric measurements (age, gender BMI, stool sample type etc.). Stool shotgun sequencing was performed for all samples.

FINRISK is a population survey that has been performed every 5 years since 1972^42^. The FINRISK 2002 survey was based on a stratified random sample of the population aged 25– 74 years from six geographical areas of Finland^43^. Stool shotgun sequencing was performed and after quality control n = 7211 participants remained available for analyses. The study protocol of FINRISK 2002 was approved by the Coordinating Ethical Committee of the Helsinki and Uusimaa Hospital District (Ref. 558/E3/2001). All participants signed an informed consent.

TwinsUK shotgun metagenomic sequencing data of human fecal samples was derived from the publicly available data from Chen et al^31^. The samples and clinical indexes were collected by Prof. Spector’s group at the King’s College London University. Participants with a history of chronic serious infection, malignant cancer or had received antibiotic treatment within 1 month before samples collection were excluded.

### Inclusion criteria

OA was defined with the help of ICD10 codes as M15-19, M16 was used for hip OA, M17 for knee OA, M18 for OA of the first carpometacarpal joint and M19 for other and unspecified OA. This was used for the FINRISK 2002 cohort, Estonia Biobank and the TwinsUK study, whereas for the Lifelines a self-assessment from a questionnaire was used.

### Exclusion criteria

All cohorts excluded participants with recent antibiotic use from microbiome collection. Healthy participants were selected by exclusion of participants with an OA or RA diagnosis. Participants with conditions known to affect gut microbiome, namely pregnancy, inflammatory bowel disease (K50-51), inflammatory bowel syndrome (K58), Celiac Disease (K90), Type I Diabetes (E10), Type II Diabetes (E11), Depression (F32-33 with recent 12 month drug prescription). After exclusion 162 OA participants remained in the FINRISK 2002 cohort, 482 in Estonia Cohort, 431 in Lifelines, and 28 in TwinsUK study.

For testing the effect of age within the FINRISK 2002 cohort participants above the age of 55 were excluded for the “<55” group and participants below the age of 65 were excluded for the “>65” group. A total of 150 participants per group were selected to resemble the number of participants in the OA analysis.

### Control matching

Next to the stringent exclusion criteria, we implemented strict control matching for each cohort, to ensure that we limit the main confounding factors. Therein, for each case (OA positive), we selected a matching control (without replacement, so every patient can only be present once in the controls) following the criteria: (i) Gender: Exact match; (ii) Age: Exact match, otherwise extend to ±1. (iii) BMI, Range of ±1, otherwise extend to ±2 etc. until ±5 (iv) Activity Score was assessed for Estonia and Lifelines cohort based on their self-assessment. These variables were considered in a custom script to look for the best possible control. If a perfect match was found (same cohort,same gender, same age, same BMI, same activity score), we chose this, otherwise we relaxed the requirements in age, BMI and activity score in a stepwise manner up to a maximum of ±5. In the end, we ensured that the parameters were not significantly different between control and cohort.

### Quality control and filtering

The quality of the reads was assessed using FastQC v0.11.9. We then trimmed the reads using kneaddata-0.12.0 with the --trf flag and otherwise standard parameters to remove the adaptor, low-quality sequences and shorts reads. Additionally, we filtered out all reads belonging to human hosts or laboratory-associated sequences.

### Taxonomic profiling

Whole genome sequencing (WGS) reads were available and used from all cohorts. We independently created taxonomic profiles for each sample with two different taxonomic profilers, analyzing all WGS samples: We processed the reads with mOTUs v 3.0.3^44^ which is based on multiple marker genes. We furthermore analyzed the same samples with the 16S rRNA based mapseq v 2.1.1 which includes hierarchical Operational Taxonomic Units (OTUs) from 90 to 99% granularity. In mapseq, we used the -skippairidcheck -fastq -paired parameters. We then merged all samples within one cohort with the -otutable and -ti 1 flag and -tl [0-4], to obtain a taxonomic profile for all different similarity thresholds (90, 96, 97, 98, 99). In mOTUs, we created an OTU-table with the merge function. An example script to profile a sample with all mentioned steps is provided in github.

### Association testing and diversity indices

The OTU-tables of both mapseq and mOTUs were then imported into R and processed further. The alpha diversity indices (Simpson and Shannon) were calculated using the vegan package version 2.6-4. To test for differences in between the alpha diversity values of case and control, we used a two-sided Welch T test with a significance threshold of p < 0.05. The ordinate method of phyloseq v1.42.0 was run to estimate the ß-diversity and clustering of the samples. We used the Bray-Curtis dissimilarity as a distance metric and the principal component analysis (PCoA) method.

For further steps, we normalized the data by calculating the relative abundances. All OTUs with a prevalence of <5% or a maximum abundance of <0.0001% were excluded. The associations were tested with a Wilcoxon Rank-sum test, adjusted for multiple testing via the Benjamini Hochberg method. P=0.05 was used as our significance threshold.

We then created a siamcat object in SIAMCAT 2.2.0 to plot the results in an association plot, and to assess fold change of OTUs in OA positive participants versus controls. The confounding factors (age, BMI, gender) were assessed in SIAMCAT via the confounding.plot function.

### Machine Learning

We used SIAMCAT 2.2.0 for machine learning and feature identification. We first normalized all features with the log.std. method. We then split our data into a 10-fold Cross validation (create.data.split (sc.obj, num.folds = 10, num.resample = 10)). We split the dataset into 10 parts and then train a model on 9 of these parts and use the left-out part to test the model. The whole process was repeated 10 times. We then trained a LASSO logistic regression classifier on our data to distinguish OA microbiota from controls. We estimated the performance on the left-out part of the data by assessment of precision and recall as well as in a receiver operating characteristic curve (ROC) analysis. Finally, we plotted the characteristics of the models (i.e. model coefficients or feature importance) with the model.interpretation.plot command.

### Enzyme-linked immunosorbent assays

LBP (serum diluted 1:500), FABP2(serum diluted 1:4), CD14(serum diluted 1:15), and S100A8/S100A9 Heterodimer complex (serum diluted 1:10) were measured from patient serum by sandwich enzyme-linked immunosorbent assay (ELISA) (LBP cat. DY870; FABP2 cat. DY3078; CD14 cat. DY883; S100A8/S100A9 cat. DY8226, R&D Systems, USA), following the manufacturer instructions. HRP conjugate was detected with enhanced chemiluminescence (ECL) substrate (cat. 32106, Pierce, USA) and measured with a microplate reader(BMG Pherastar FS, BMG LABTECH, Germany).

Zonulin (serum diluted 1:10) were measured from patient serum by sandwich enzyme-linked immunosorbent assay (ELISA) (cat. E-EL-H5560, Elabscience, USA), according to the manufacturer instructions. Optical density was measured with the microplate reader (BMG Pherastar FS, BMG LABTECH, Germany) set to read the absorbance at 450 nm with a wavelength correction set at 540 nm.

All ELISA assays were performed in High-Throughput Screening (HTS) mode with reagent and liquid dispensing by dispenser (CERTUS FLEX Fritz Gyger AG, Germany), serum sample dispensing by dispenser (ECHO 650, Labcyte, US) or manually, and washes by plater washer (EL406, Agilent BioTek, USA) as previously described^45^. All HTS assays were performed at the Institute for Molecular Medicine Finland (FIMM) High Throughput Biomedicine unit. Statistical differences between groups were calculated using a two-sided t-test and considered significant with p <0.05.

### Protein catalogue

A microbial function association was inferred by mapping the metagenomes of the cohorts to a protein catalogue of the gut microbiome provided by the Unified Human Gastrointestinal Genome (UHGG) version 2.0.1^46^. The UHGG is available for download via MGnify^47^. The UHGG represents a snapshot of publicly available data of the human gut microbiome, including 13,910,025 representative proteins (referred to as UHGP-90), which represent a clustering of all proteins at the 90% amino acid identity. In addition, the UHGP-90 proteins were annotated with eggNOG, InterPro, COG and KEGG.

The original protein clustering was conducted using the ‘linclust’ algorithm from the MMseq2 package^48^. The ‘linclust’ algorithm is an ultra-fast clustering algorithm, capable of clustering hundreds of millions of proteins. However, our preliminary investigation revealed that a significant proportion of representative proteins exhibited an amido-acid identity exceeding 90%. The UHGP-90 dataset was re-clustered with a more sensitive clustering algorithm implemented in CD-HIT version 4.8.1^49^ with the parameters -c 0.9 -G 1 -aL 0.9 -aS 0.9 -g 1. This resulted in the almost complete removal of protein pairs exhibiting an amido acid identity exceeding 90%. The number of proteins in the UHGP-90 was reduced by 11%, leaving 12,383,493 representative proteins (referred to as UHGP-90’) with at least 10% amino-acid identity divergences.

### Mapping

The metagenomes of the patients were subjected to quality and adapter trimming using the wrapper script trim_galore (FelixKrueger/TrimGalore: v0.6.7) with the parameters --paired -- length 90. To eliminate reads derived from the host genome, the libraries were aligned with Bowtie2^50^ using the default parameters to the human reference genome GRCh38. Only reads with no alignment were retained. The quality of the libraries was evaluated prior to and following the cleaning process with FastQC.

The cleaned libraries were mapped to representative proteins UHGP-90’ using a translated DNA search with the software diamond and the mode ‘blastx’^51^. The parameters --min-score and --min-orf were adapted to the read lengths to enforce essentially global alignments on the read, while being more permissive on the alignment score. For the TwinsUK cohort, the parameters were set to ‘--fast --min-score 50 --min-orf 30’, while for the EstBB cohort, they were set to ‘--fast --min-score 75 --min-orf 45’. In addition, the parameter ‘–-top 5’ was introduced to collect the highest-scoring matches and those with marginally lower scores. Subsequently, the matches were post-processed to filter for highly specific mappings by removing reads mapping to more than 4 proteins.. The UHGP-90 provides a KEGG annotation for 39% of representative proteins. The majority of these are associated with a single KEGG orthologue. In contrast, only 4% of the proteins are annotated for Gene Ontology (GO), but if a protein is annotated, then a considerable number of GO terms are associated with it. The present study focuses on the analysis of the KEGG annotation, as the KEGG annotation covers a significantly larger number of proteins. However, the findings obtained using the GO annotation do not deviate from the reported results.

Firstly, the number of counts per representative protein was calculated. A count of one was attributed to each read, distributed equally among all the proteins that it was mapping to (max. 4 proteins). Subsequently, the number of counts per KO was calculated as the sum of the protein counts associated with a given KO.

The mapped read counts were merged into a single matrix containing all patients versus all proteins. Only proteins with more than 10 hits in at least 10 participants were retained. This filtered count matrix was normalised and log transformed by adding a pseudo count of 10 to each number, dividing each number by the total count number of each library (library size normalisation) and by log transformation.

### WGCNA

A Weighted Gene Correlation Network Analysis (WGCNA) was performed on the normalized counts of participants versus KO gene families using the R package WGCNA^52^. The co-abundances were constructed by calculating signed adjacency matrices using a soft-thresholding power of 6 and a pairwise Pearson correlation between all genes. Subsequently, a signed topological overlap matrix (TOM) was calculated from each adjacency matrix, converted to distances, and clustered by hierarchical clustering using average linkage clustering. The modules were identified in the resulting dendrogram using the Dynamic Hybrid tree cut with a cut height of 0.995 and a minimum module size of 20 genes. These settings produce relatively fine-grained and signed modules, referred to as modules eigengenes (ME). The TwinsUK cohort resulted in 30 such MEs, while the EstBB cohort resulted in 37 MEs (Figure 5). The correlation values of the MEs were tested for statistical significance using the function corPvalueStudent of the R package WGCNA.

The ME cluster has tens to hundreds (up to 2030) of KO. To define gene sets of interest within the MEs, a measure V_(i,k) relating gene i and trait k over all modules by

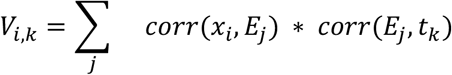

where corr() is the Pearson correlation; *xi* is the abundance vector of gene *i, E*_*j*_ the ME of module *j* and *t*_*k*_ the trait value, either binary or continuous.

Given 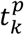 the *p*^*th*^ random permutation of the trait *t*_*k*_, one can compute Z-score *Z*_*i,k*_ as

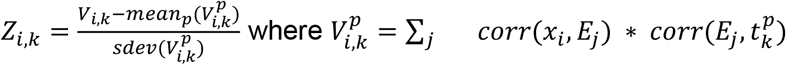

over 10’000 permutations. Z-score with absolute value equal or greater than 3 were considered significant and used to define significant gene sets.

A functional enrichment analysis using Gene Set Enrichment Analysis (GSEA) was conducted on the aforementioned gene sets with the function enrichKEGG of the R package clusterProfiler version 4.8.3^53^. The background gene set (universe) comprised all KO in the normalised abundance matrix. Statistical significance was adjusted for multiple testing using the Benjamini-Hochberg procedure (BH). Enriched pathways were retained for an adjusted p-value of ≤0.01.

### Statistical analysis

All statistical analyses were performed in R.

## 3. Results

### Study cohorts used in this work

We analyzed the gut microbiota of 1395 OA patients from 4 European cohorts, namely the FINRISK 2002 cohort, Estonia microbiome cohort (EstMB) from the Estonian Biobank, the Lifelines and the TwinsUK study. There were a total of 191 prevalent OA cases from a total of 8000 participants in the FINRISK 2002 cohort, 590 out of 7019 in Lifelines, 557 out of 2509 in the EstMB cohort and 57 out of 250 from the TwinsUK study. 20 of the OA patients from the TwinsUK cohort were twins. Across cohorts OA participants were predominantly female (72% ± 12%) with a mean age of 60,4 ± 4,7 and BMI 27,6 ± 1,7 kg/m^2^ (Table1). After applying stringent exclusion criteria (see methods), we excluded a total of 292 cases. We then matched the OA cases with healthy controls which led to further exclusion of 136 unmatched cases. Healthy controls were selected from a pool of total 17 ‘641 participants from all four cohorts defined by not being diagnosed with OA or RA to whom the same exclusion criteria were applied. We analyzed the metagenomic samples of each cohort and uniformly processed sequencing and phenotypic data with the same pipelines to ensure comparability across the different studies. The selected cases were analyzed for microbiome composition for all cohorts, functional gene analysis for EstMB and TwinsUK cohort and serum biomarker analysis for FINRISK samples (Figure 1).

**Table 1.** Participant information

|  | EstMB | Lifelines | FINRISK | TwinsUK | All |
| --- | --- | --- | --- | --- | --- |
| **# Prevalent OA cases** | 557 | 590 | 191 | 57 | 1395 |
| **# OA cases after exclusion** | 482 | 431 | 162 | 28 | 1103 |
| **# OA cases after matching** | 346 | 431 | 162 | 28 | 967 |
| **Mean Age** | 54.8 | 57.01 | 65.5 | 65 | 60.4 |
| **Mean BMI [kg/m<sup>2</sup>]** | 27.75 | 26.99 | 30.28 | 25.9 | 27,6 |
| **Female participants [%]** | 69.98% | 66.44% | 57.89% | 100% | 72% |

**Figure 1.**
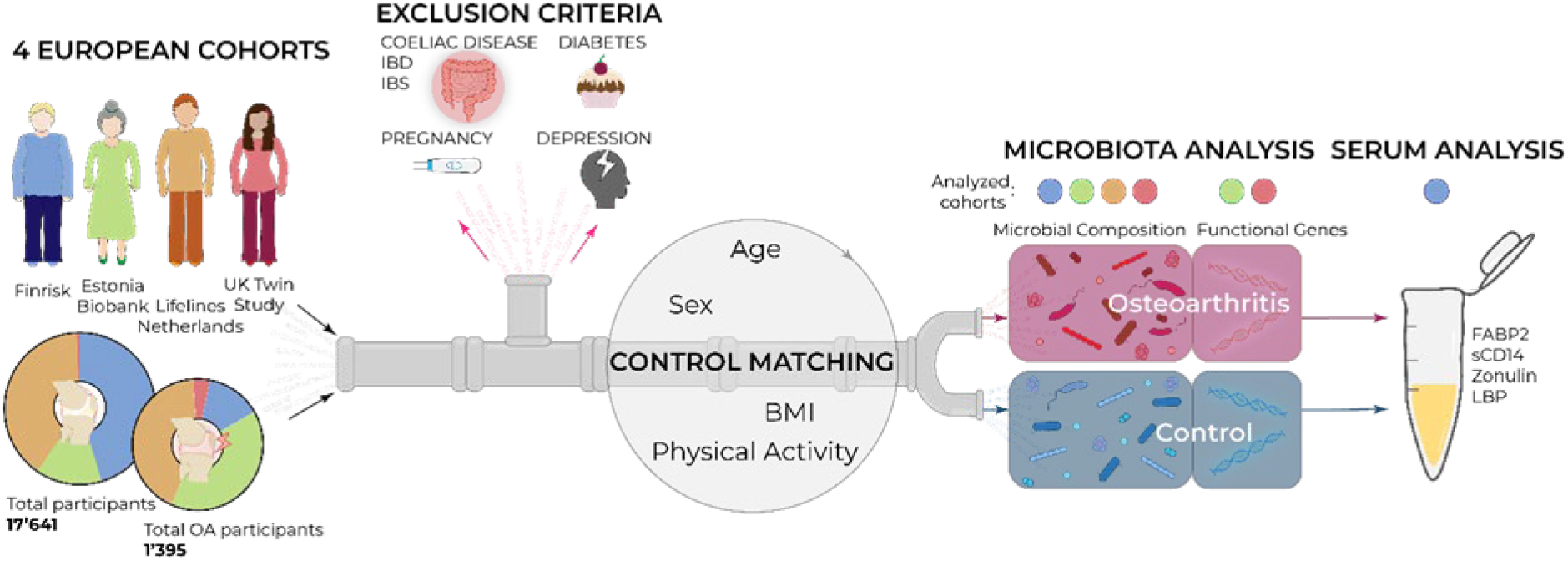
Multi-cohort analysis overview, FINRISK 2002 (blue), EstMB (green), Lifelines (orange), TwinsUK (red).

### No differences in microbial diversity

We first explored whether the gut microbiome of patients with OA differs in species richness or composition from controls. We therefore calculated α-diversity indices (Figure 2a, Figure S1) for all cohorts. We observed no significant differences (p>0.05) between cases and controls in any of the four cohorts.

**Figure 2.**
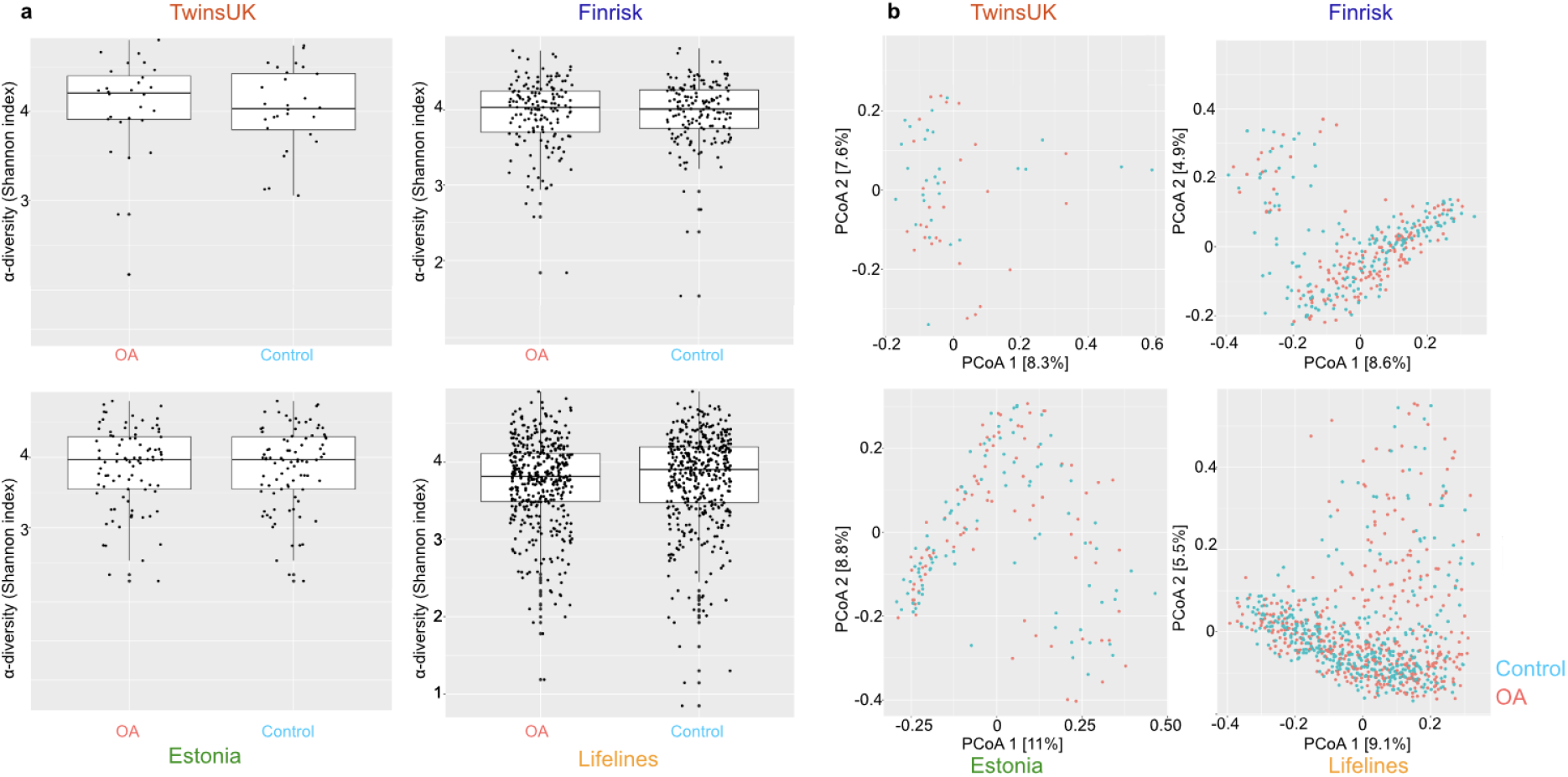
Microbial diversity in the gut microbiome of OA patients of all four different cohorts. a) α-diversities represented by Shannon index. Each dot corresponds to a different alpha-diversity value. Boxes represent the first and third quartiles (25% and 75%), the inner horizontal line the median. b) β-diversity (Bray Curtis) represented with PCoA in each of the four cohorts comparing OA (red) and control (blue) cases.

Additionally, we estimated the ß-diversity of all samples with Bray-Curtis dissimilarity and visualized the grouping using principal component analysis (PCoA). Also here, no separation of the microbiota of OA patients was visible in comparison to the healthy controls in any of the four studied cohorts (Figure 2b). While there were some microbiome clusters, these did not correspond to OA, age, BMI or sex (Figure S2).

### No single taxon is significantly enriched in Osteoarthritis patients

Due to the compositional nature of microbiomes, we next normalized our data by calculating the relative abundances and performed a differential abundance analysis according to state-of-the art methods^54^. Remarkably, not a single operational taxonomic unit (OTU) classified with mOTUs was enriched significantly in the FINRISK 2002 cohort (Figure 3a). We verified this observation in the other three cohorts, and none of them contained a significantly differential abundant OTU (Figure S3). While some species appear more prevalent in OA patients in the FINRISK 2002 cohort, such as *Roseburia intestinalis* and *Clostridium sp*, neither of them reached statistical significance and these trends were not repeated in all the other cohorts (Figure 3a, Figure S3). We further hypothesized that changes in microbiome compositions could occur also at a higher or lower phylogenetic level - e.g. that while there are no species-level differences estimated by mOTUs, a microbial genus or subspecies is enriched in either group. Hence, we used the 16s rRNA marker gene based profiling tool mapseq^55^ to verify and extend the results, on different OTU similarity thresholds ranging from 90% (approximately family level) to 99% (subspecies level). With both mapseq or mOTU approaches we did not find any taxonomic group to be significantly enriched (Figure 3a, Figure S3, S4).

**Figure 3.**
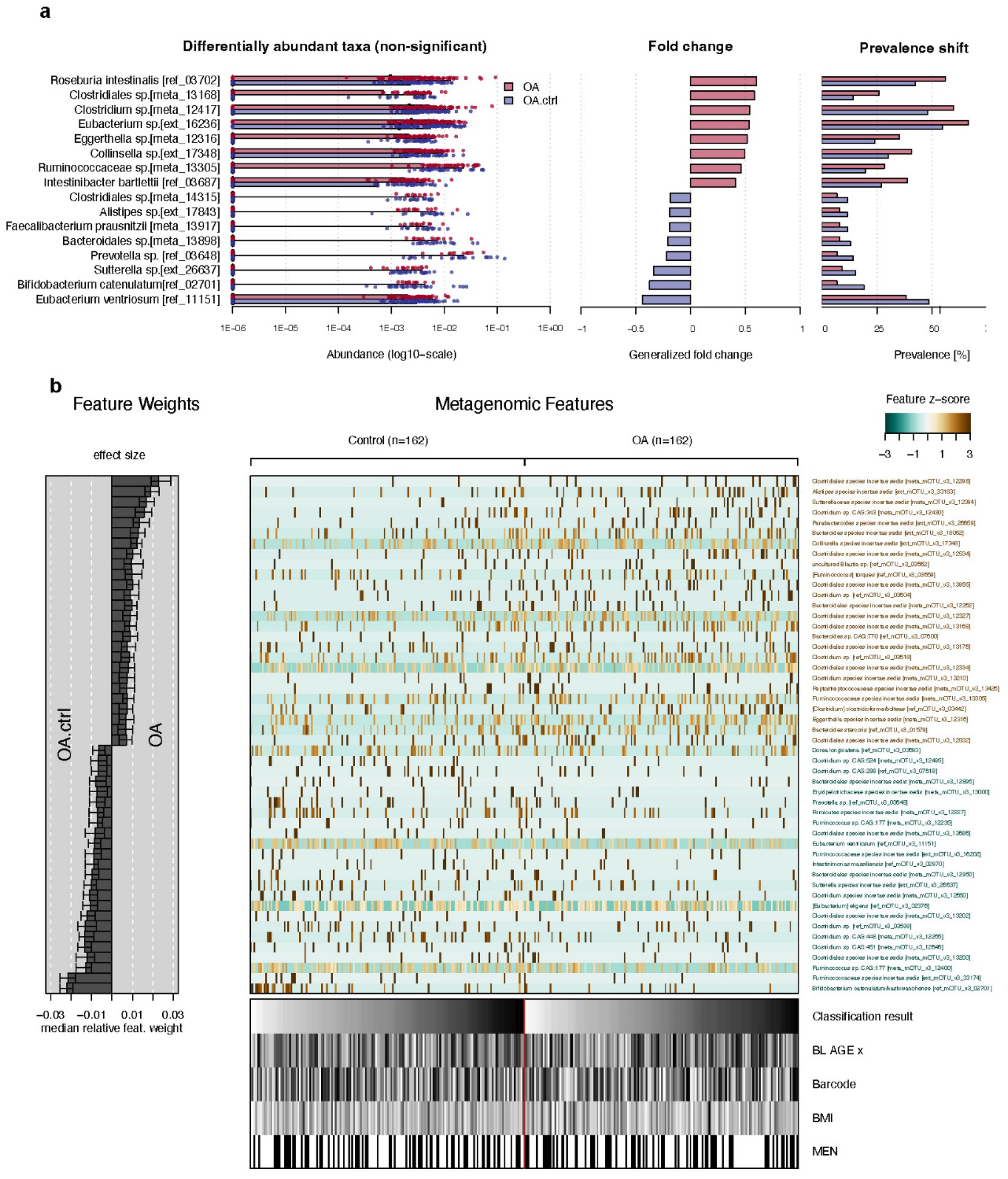
Assessment of enriched taxa in the gut microbiota of OA participants in FINRISK 2002 cohort a) 8 differentially abundant taxa (non significant) enriched in OA (red) and control (blue), as well as the associated differences in foldchange (middle panel) and prevalence (right panel). b) Least absolute shrinkage and selection operator (LASSO) model for FINRISK 2002, showing the most important feature weights (in this case, OTUs). On the bottom, meta variables (Age, BMI and gender) are shown.

To find more complex trends that may involve a combination of OTUs recognizable by a machine learning model, we built a least absolute shrinkage and selection operator (LASSO) model for each cohort within SIAMCAT 2.2.0^54^. We implemented this model in order to make sure we can pick up and interpret all the features even if they are not visible with simple association testing. However, this analytical approach also showed no significant differences between OA and the controls since the model was unable to perform better than classifying by chance (AUROC=0.554, Figure 3b).The model was verified by using age categories for the same data (Figure 4).

**Figure 4.**
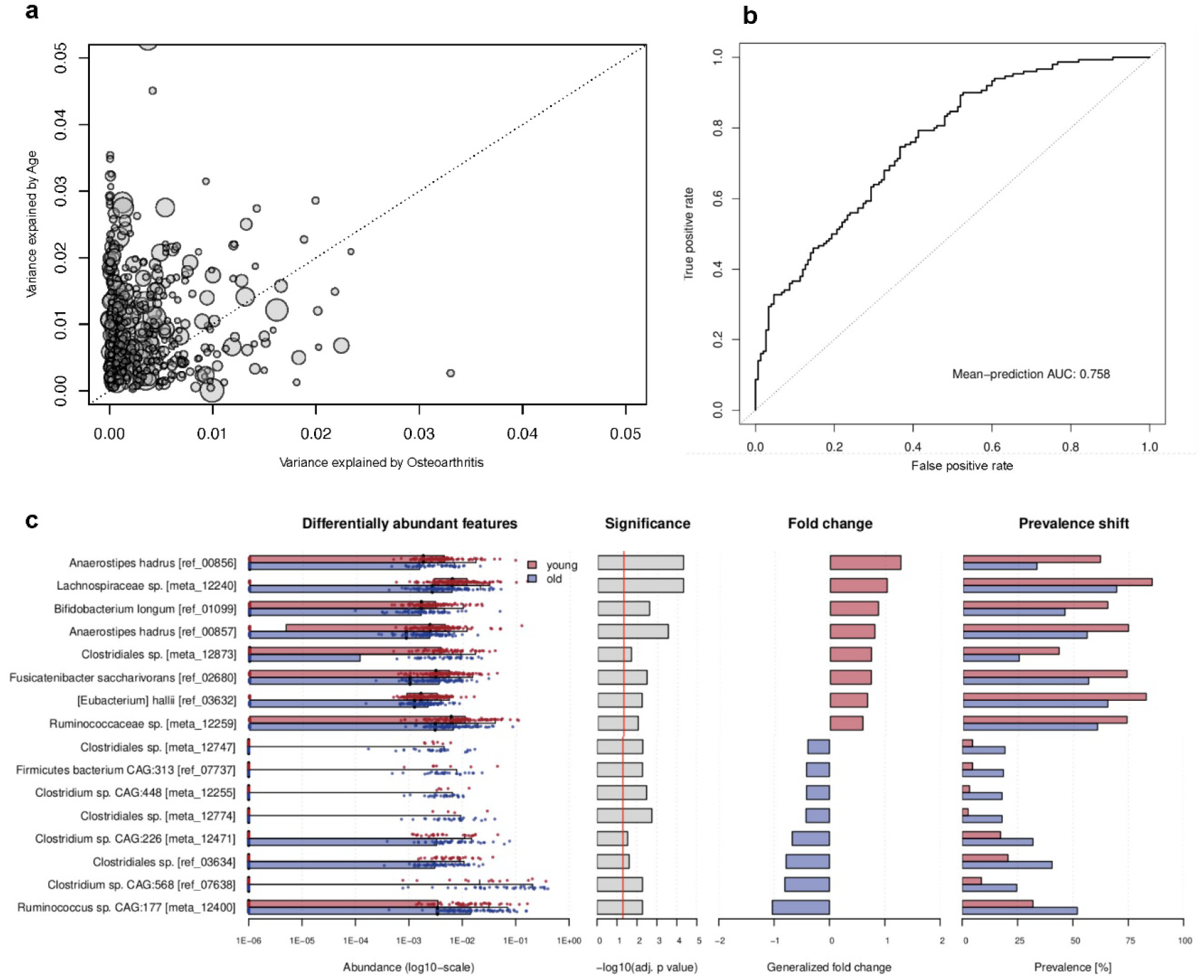
Effect of age on microbial composition in FINRISK 2002 cohort a) Variance explained by age and OA. Each dot corresponds to one participant. b) AUROC Curve of the LASSo model calculated for the FINRISK 2002 model to distinguish between old (>65 years) and young (<55 years) participants. c) 8 differentially abundant taxa enriched in <55 group (red) and >65 group (blue) are shown, a full list is provided in Supplementary Figure S5. On the right hand side, fold changes and prevalence shifts of the OTUs are plotted.

**Figure 5.**
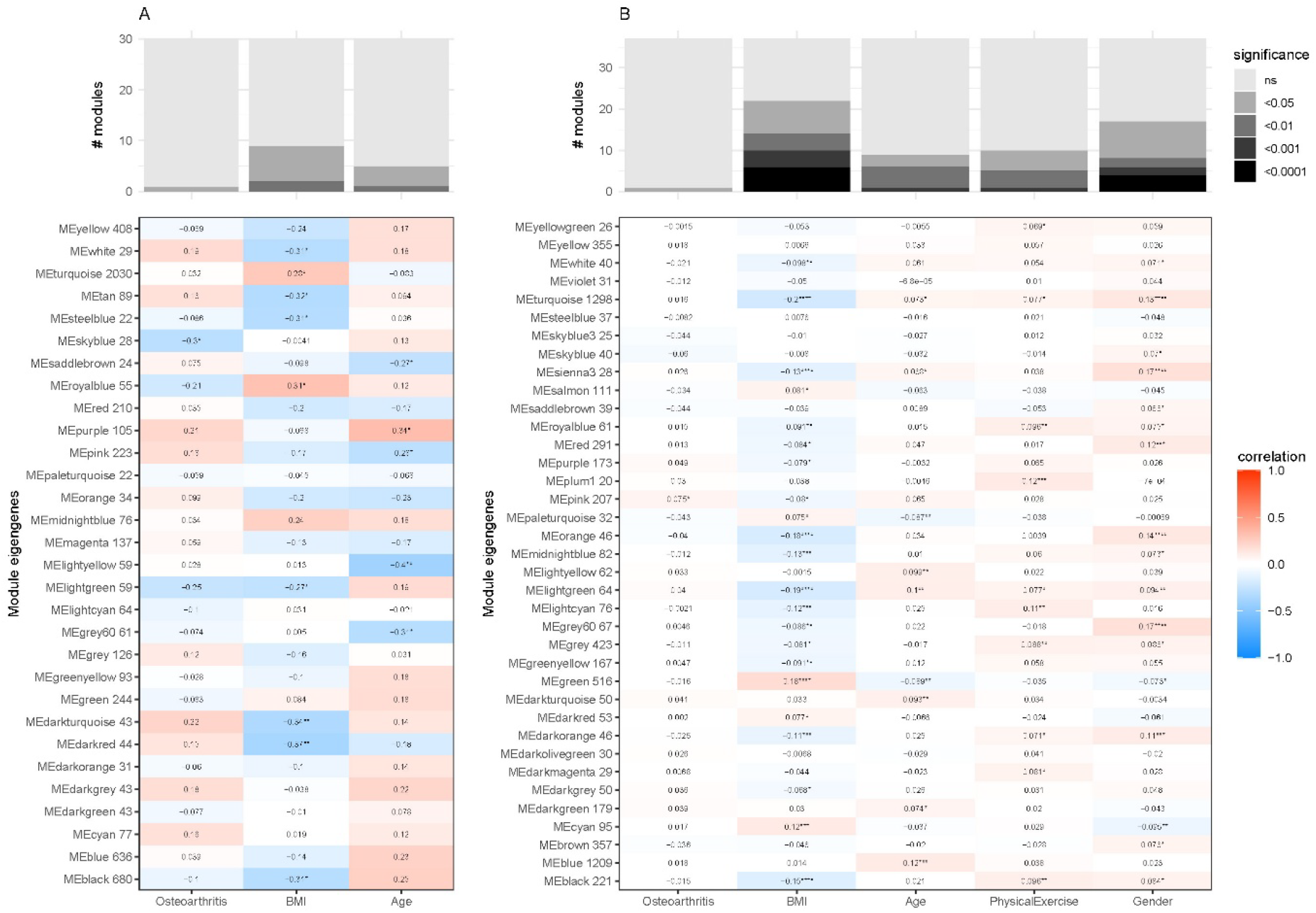
The heatmaps illustrate the Pearson correlation between eigengene (functional) modules and each trait. Figure A shows the results for the TwinsUK cohort, while Figure B shows the results for the EstBB cohort.The eigengene modules are derived from an unsupervised weighted gene co-expression network analysis (WGCNA) of read counts on KEGG orthologues. The names of the eigengene modules are followed by the number of KEGG orthologues that are clustered together. The names of the eigengene modules identified in A and B are not interrelated. The values represented in the heatmap indicate the correlation coefficient, with the stars denoting the level of statistical significance, as follows: *: p < 0.05; **: p < 0.01; ***: p < 0.001; ****: p < 0.0001. The bar plots displayed at the top illustrate the number of modules identified at each level of significance and for each trait.

### The microbiome of older patients with a higher BMI contributes more to microbial diversity than OA

Upon checking potential confounding factors by generalized linear models for the analyzed cohorts, we observed that especially age, but also BMI, both known OA-risk factors, could influence microbiome composition (Figure 4a). In the plot, the variance explained by age is plotted against the variance explained by the disease, and it is apparent that most points are on the left side, meaning that age together with associated changes in e.g. BMI influences the microbiome composition stronger than OA. To verify our findings, we repeated our primary analysis pipeline in the FINRISK 2002 cohort with two different age groups, <55 years and >65. Here, the trained LASSO model was able to distinguish between <55 and >65 participants with an AUROC of 0.78, indicating that there are differences in the microbiota (Figure 4b). Indeed, we observed significantly enriched and depleted OTUs between >65 and <55 participants (Figure 4c). A complete list of significant associations is provided in Figure S5. *Anerostipes hadrus, Lachnospiraceae sp. and Bifidobacterium longum* were among the most enriched OTUs in <55 participants. On the other hand, *Ruminococcus sp and multiple Clostridium sp*. OTUs were significantly more abundant in the >65 group.

### Weighted Gene Correlation Network Analysis reveals BMI, not OA related changes

Next to the taxonomic composition also the functional potential of the microbial community matters for the human host. We therefore searched for enriched or depleted KEGG orthologues (KO)^56^ in OA participants or healthy controls using an unsupervised Weighted Gene Correlation Network Analysis (WGCNA). This analysis revealed several KO modules (sets of KOs with similar patterns of enrichment or depletion) that were differently enriched between investigated traits. BMI appeared to be differentially enriched more often among participants than sex. KO modules related to age and physical activity (PhysicalExercise) are also differentially distributed among participants, but to a lesser extent. KO modules related to OA are the least differentially distributed among participants. The observed pattern was similar across both studied cohorts; however, the effect was more pronounced in the EstMB cohort.

Only a single pathway was found to be significantly enriched with one of the traits. The pathway ko02040 ‘Flagellar assembly’ is significantly enriched (p.adj=1e-07) for BMI in the TwinsUK cohort.

The correlation between trait and abundance of the KOs was, on average, significantly higher in the TwinsUK cohort than in the EstMB cohort. However, despite this, most correlation coefficients are not significant. This discrepancy is likely due to the differing sample sizes of the two cohorts.

### Gut permeability biomarkers are elevated in older participants but not in OA

As the gut-joint axis implication in OA does not solely depend on gut dysbiosis we also assessed intestinal permeability and the degree of systemic endotoxemia by measuring the concentration of soluble CD14, FABP2, LBP and Zonulin in the serum of the selected OA and matched control participants from the FINRISK 2002 cohort. No significant differences in these biomarkers were observed between OA patients and healthy controls (Figure 6a, c, e, g). Next, we compared the serum levels of these biomarkers in the “<55” and “>65” age groups. In line with our previous results, we observed a significantly higher concentration of sCD14, FABP2 and Zonulin in the serum of older participants in the “>65” group (Figure 6b, d, h).

**Figure 6.**
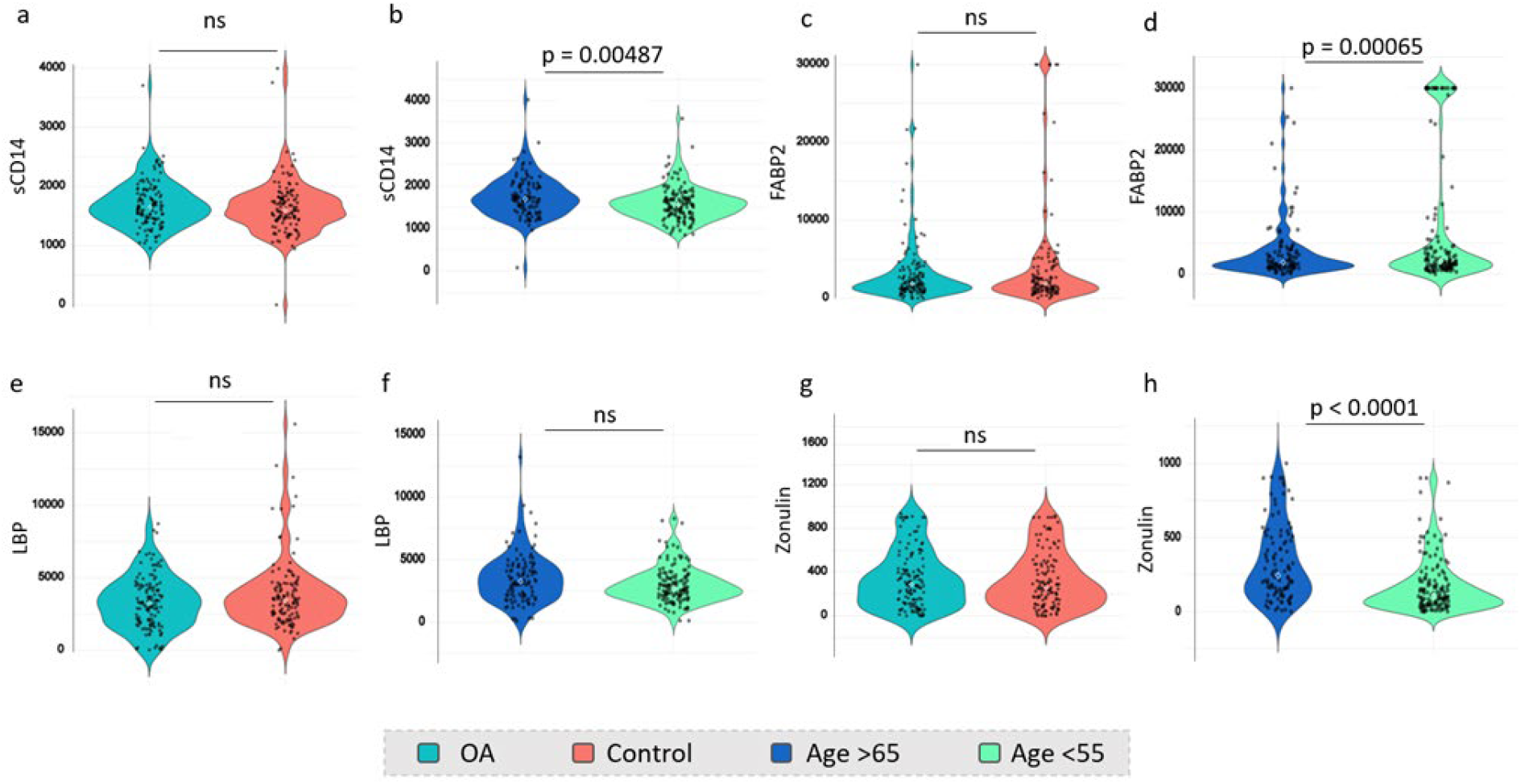
Serum intestinal permeability biomarker analysis, concentration of a-b) sCD14 c-d) FABP2 e-f) LBP g-h) Zonulin in OA (aquamarine) compared to control participants (red) and in “>65” age group (dark blue) compared to “<55” age group (green) participants. Statistical significance was determined via a Welch Two Sample t-test.

## 3. Discussion

We observed no significant difference in species and community composition diversity (alpha and beta diversity) between OA and control groups (Figure 2a, b). In the PCoA analysis we found different microbial clusters in different cohorts. These clusters do not correspond to OA or any OA risk factors (Figure S2). It has previously been described that distinct microbial enterotypes correspond to differences in microbial composition that cannot be explained by age, BMI, sex, geographical location^57^. We therefore believe that the clusters could reflect the different enterotypes of the gut microbiome or differences in diet^5,57– 61^.

Furthermore, we were not able to identify any statistically significantly enriched microbial species in OA patient microbiota (Figure 3a). In concordance, our LASSO model could not accurately predict whether the microbiota belonged to an OA patient or not (Figure 3b). This means that even more complex combination patterns of microbial species abundances, which can be missed in the association analysis, were not enough to distinguish cases from controls. This did not change when stratifying based on affected joint (Figure S6). In line with previous literature^35,57,62^we observed that age was the key factor significantly impacting the microbial gut community (Figure 4a). Interestingly, multiple *Clostridium sp*. OTUs were significantly more abundant in the “>65” group (Figure 4c). The connection between Clostridia and advanced age has already been reported^9^. This could explain why some studies previously observed higher Clostridia abundance in OA patients as age is a risk factor for OA^29^. Furthermore, both Clostridiales and *Ruminococcus sp*., found abundant in the >65 group, belong to the Firmicutes phylum, an increase of which is associated with bad gut health and obesity^63,64^. The higher abundance of these bacteria could also influence the Firmicutes/Bacteroides ratio that has been reported to be changed in the OA gut microbiome as well as in the aging population^28,29,31^. Interestingly, *Anaerostipes hadrus* as well as *Lachnospiraceae sp* were found more abundant in <55 participants. Both bacterial species are butyrate-producing bacteria associated with good gut health and increased expression of epithelial tight junctions^65,66^. This data again suggests that some of the bacterial changes reported to be associated with OA can be attributed to age.

We also looked at the differences in bacterial genes present in gut microbiota of OA participants compared to controls and found that there are no OA associated KO terms (Figure 5). We did observe some high BMI-related features such as enriched flagellar assembly (Figure 5). This is a good control for our bioinformatic approach as this pathway has previously been reported to be associated with obesity^67^. This finding that BMI has more statistically significant enriched bacterial gene features could also help explain some of the reported OA related differences in microbial composition, as high BMI is also a risk factor for OA. Overall, we observed higher contributions to changes coming from all the risk factors including age, sex and self-reported levels of physical activity. This suggests that the risk factors have a higher impact on microbial genes than the OA disease state itself.

Furthermore, the involvement of leaky gut in OA pathogenesis does not seem to be directly linked to disease state according to serum biomarker analysis. We found no significant increase in FABP2, sCD14, zonulin nor LBP in the serum of OA participants compared to their matched healthy controls (Figure 6). However, we did observe aging related changes in sCD14, FABP2 as well as Zonulin (Figure 6). This suggests that OA itself does not cause gut leakiness, but it could be induced by aging. However, it could still be a contributing factor to disease progression, as we did not analyze pain in this study. There are previous studies linking OA with LBP, for example a recent study by Binvignat et al showing that serum LBP and zonulin related proteins are associated with erosive hand OA^20^. However, in our study we did not look at the disease clinical subtypes which may provide further insight into the connection of specific clinical symptoms in OA and gut permeability.

Together, our data suggests that in the analyzed cohorts, no significant association exists between the gut microbiome and OA disease state, and that previously reported differences likely stem from lack of exclusion criteria, small sample size and/or lack of age and BMI matching. In conclusion, OA patients likely have a slightly different microbiome composition compared to the general population; however, it does not seem to be associated with the OA disease state but instead with their age, BMI and sex. Our findings highlight the importance of rigorous study design in microbiome research and challenge the hypothesis that OA is linked to gut dysbiosis. By using state-of-the-art analytical approaches and applying stringent controls, we contribute to a more precise understanding of host-microbiome interactions, underscoring the need for high-quality evidence in the exploration of microbiome-related disease mechanisms.

## ACKNOWLEDGEMENTS

The authors wish to acknowledge the services of the Lifelines Cohort Study, the contributing research centers delivering data to Lifelines, and all the study participants. As well as the Department of Twin Research of the TwinsUK study and the participants of all the cohorts.

We acknowledge the personal and the services of the Estonian Biobank for their sample collection and technical assistance. We thank Mait Metspalu, Andres Metspalu, Lili Milani and Tõnu Esko from Estonian Biobank research team for Estonian Biobank health data collection. Data analysis for the EstMB cohort was carried out in part in the High-Performance Computing Centre of University of Tartu, and we thank HPC Support Team of the Institute of Computer Science at the University of Tartu for delivering exceptional service and assistance in installing the necessary programs on the cluster.

We acknowledge the personnel at the FIMM High Throughput Biomedicine Unit, which are hosted by the University of Helsinki and supported by HiLIFE and Biocenter Finland, for their expert technical assistance.

We would also like to thank Dr. Alexander Harms and Dr. Shipin Zhang from ETH Zurich for their feedback on the manuscript drafts.

## AUTHOR CONTRIBUTIONS

KB, CVM, GB and MZW contributed to the conception and design of the study. KB, CvM, GB and MZW obtained a funding source. KB was responsible for the collection of data and data interpretation. VS, AH and TN contributed to the collection of the FINRISK 2002 data and RK carried out the shotgun sequencing of the stool samples. EO contributed to the collection of the EstMB data. LM, SN, VDT, MP, KP, MJ and KB were involved in data analysis. KB and LM wrote the draft of the article. All authors contributed to the critical revision of the article and have given their final approval of the article.

## ROLE OF THE FUNDING SOURCE

This project has received funding from the European Union’s Horizon Europe research and innovation programme under grant agreement No 101095084. This work was supported by the Swiss State Secretariat for Education, Research and Innovation (SERI) under contract number 22.00462.

The Lifelines initiative has been made possible by subsidy from the Dutch Ministry of Health, Welfare and Sport, the Dutch Ministry of Economic Affairs, the University Medical Center Groningen (UMCG), Groningen University and the Provinces in the North of the Netherlands (Drenthe, Friesland, Groningen).

EstMB is funded by the Estonian Center of Genomics/Roadmap II project No 16-0125, by Estonian Research Council grant PRG1414 and EMBO Installation grant No 3573.

TwinsUK is funded by the Wellcome Trust, Medical Research Council, Versus Arthritis, European Union Horizon 2020, Chronic Disease Research Foundation (CDRF), Wellcome Leap Dynamic Resilience Programme (co-funded by Temasek Trust), Zoe Ltd, the National Institute for Health and Care Research (NIHR) Clinical Research Network (CRN) and Biomedical Research Centre based at Guy’s and St Thomas’ NHS Foundation Trust in partnership with King’s College London.

## Data availability

Because the data comes from samples and information of human participants, due to the required data protection agreements between the participants and the used biobanks as well as European law, the individual-level data is not available. Data may be obtained from a third party and are not publicly available. Researchers can apply to the respective cohorts to access the cohort data used in this study. More information about how to request the data and the conditions of use can be found on their respective websites.

### Code availability

All custom code used in this analysis to reproduce the results is uploaded on github: https://github.com/lukasmalfi/osteoarthritis

