## Supplementary Information for "Microbiome variations in osteoarthritis reflect aging and metabolic factors, not the disease"

**Extended data**

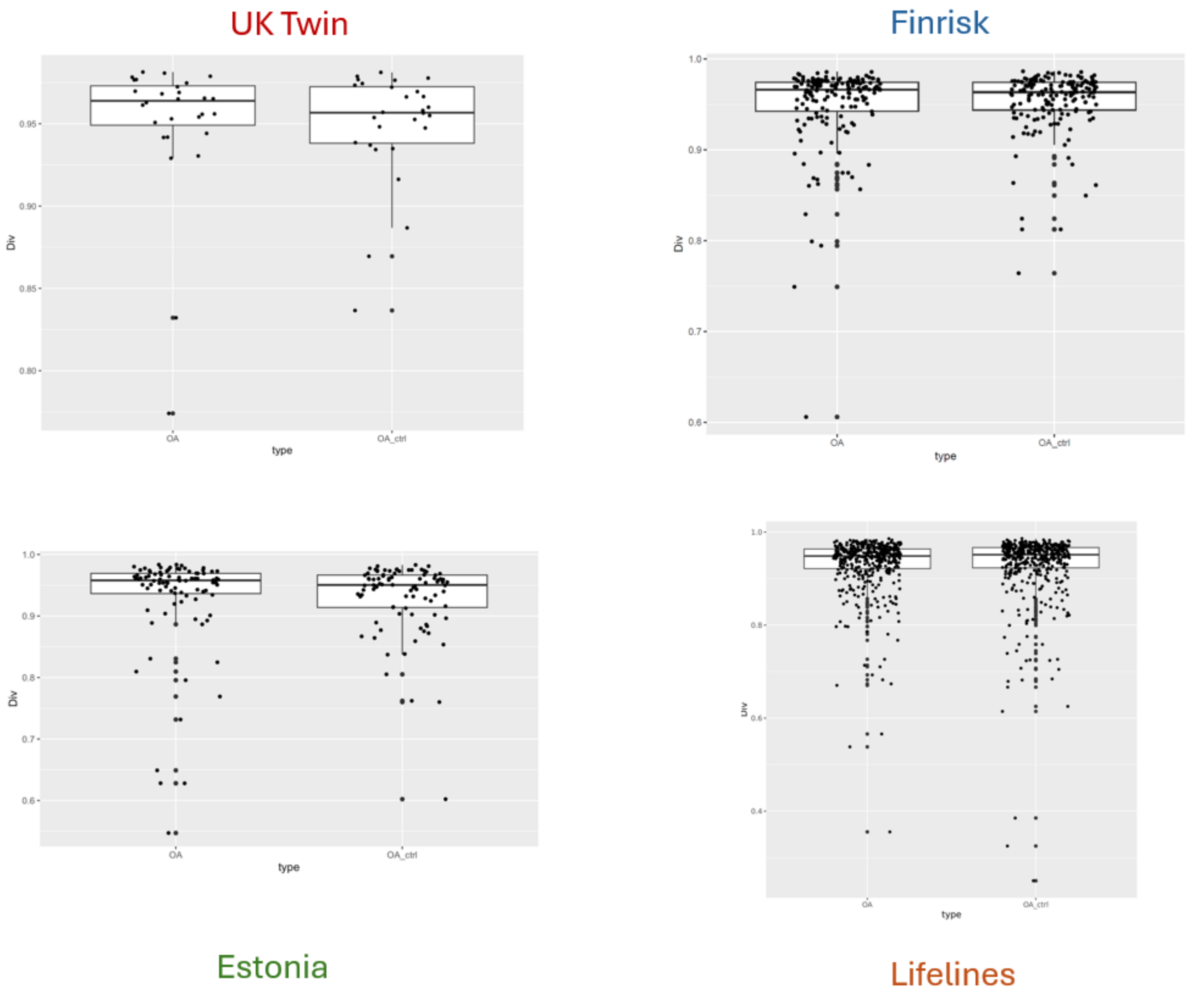


**Supplementary Figure 1** Microbial diversity in the gut microbiome of OA patients α-diversity represented by Simpson index

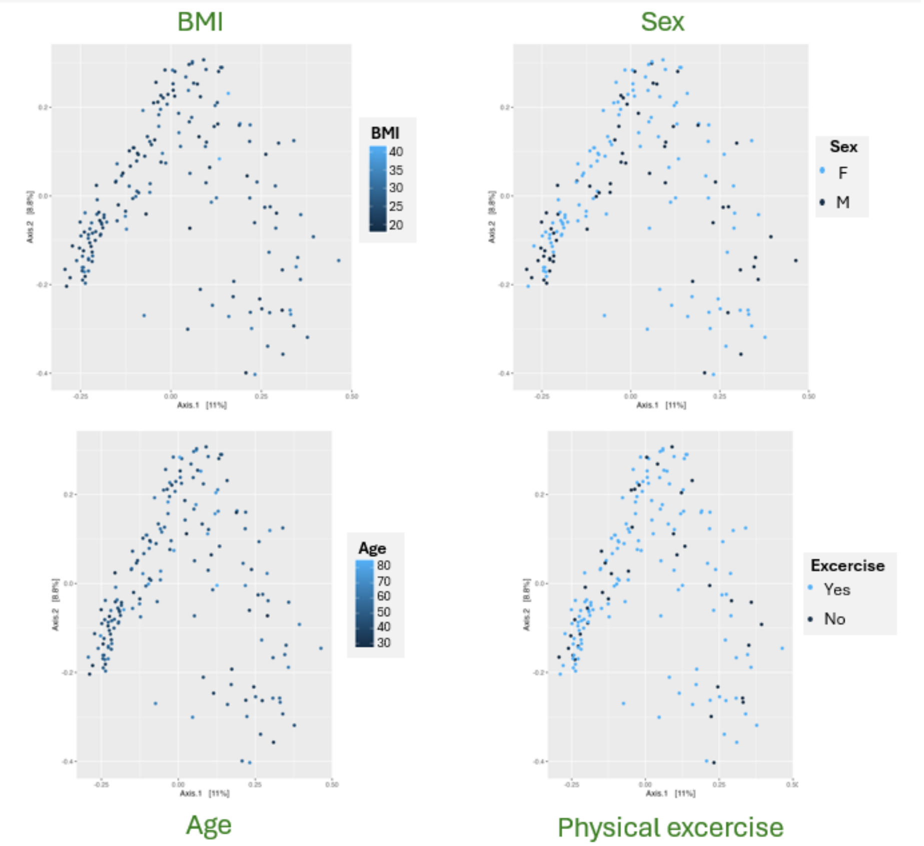


**Supplementary Figure 2** PCOA of Bray Curtis colored due to age, sex and BMI represented by the Estonia cohort

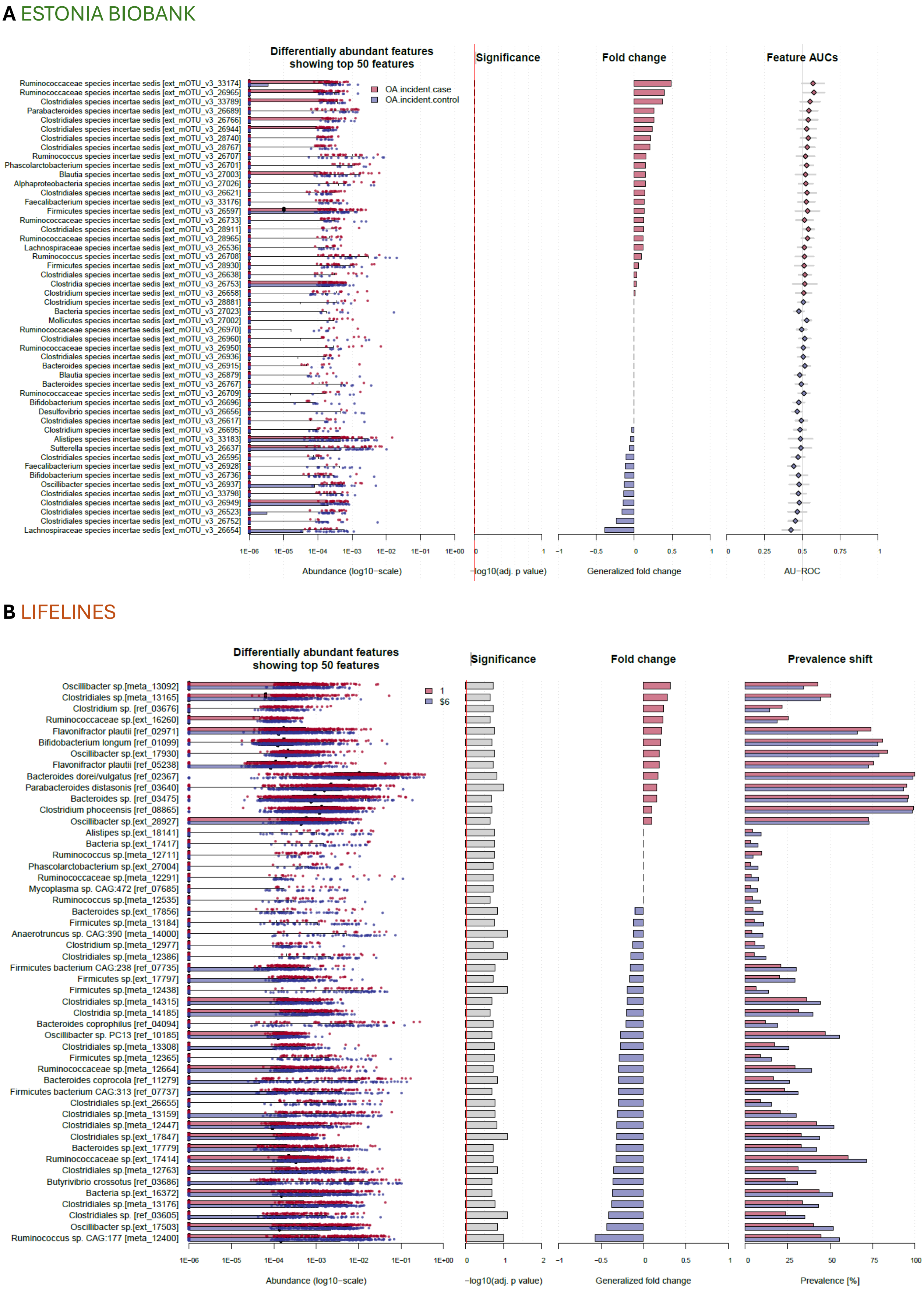
A ESTONIA BIOBANK
Differentially abundant features showing top 50 features
Significance
Fold change
Feature AUCs
OA.incident.case
OA.incident.control
Ruminococcaceae species incertae sedis [ext_mOTU_v3_33174]
Ruminococcaceae species incertae sedis [ext_mOTU_v3_26965]
Clostridiales species incertae sedis [ext_mOTU_v3_33789]
Parabacteroides species incertae sedis [ext_mOTU_v3_26689]
Clostridiales species incertae sedis [ext_mOTU_v3_26766]
Clostridiales species incertae sedis [ext_mOTU_v3_26944]
Clostridiales species incertae sedis [ext_mOTU_v3_28740]
Clostridiales species incertae sedis [ext_mOTU_v3_28767]
Ruminococcus species incertae sedis [ext_mOTU_v3_26707]
Phascolarctobacterium species incertae sedis [ext_mOTU_v3_26701]
Blautia species incertae sedis [ext_mOTU_v3_27003]
Alphaproteobacteria species incertae sedis [ext_mOTU_v3_27026]
Clostridiales species incertae sedis [ext_mOTU_v3_26621]
Faecalibacterium species incertae sedis [ext_mOTU_v3_33176]
Firmicutes species incertae sedis [ext_mOTU_v3_26597]
Ruminococcaceae species incertae sedis [ext_mOTU_v3_26733]
Clostridiales species incertae sedis [ext_mOTU_v3_28911]
Ruminococcaceae species incertae sedis [ext_mOTU_v3_28965]
Lachnospiraceae species incertae sedis [ext_mOTU_v3_26536]
Ruminococcus species incertae sedis [ext_mOTU_v3_26708]
Firmicutes species incertae sedis [ext_mOTU_v3_28930]
Clostridiales species incertae sedis [ext_mOTU_v3_26638]
Clostridia species incertae sedis [ext_mOTU_v3_26753]
Clostridium species incertae sedis [ext_mOTU_v3_26658]
Clostridium species incertae sedis [ext_mOTU_v3_28881]
Bacteria species incertae sedis [ext_mOTU_v3_27023]
Mollicutes species incertae sedis [ext_mOTU_v3_27002]
Ruminococcaceae species incertae sedis [ext_mOTU_v3_26970]
Clostridiales species incertae sedis [ext_mOTU_v3_26960]
Ruminococcaceae species incertae sedis [ext_mOTU_v3_26950]
Clostridiales species incertae sedis [ext_mOTU_v3_26936]
Bacteroides species incertae sedis [ext_mOTU_v3_26915]
Blautia species incertae sedis [ext_mOTU_v3_26879]
Bacteroides species incertae sedis [ext_mOTU_v3_26767]
Ruminococcaceae species incertae sedis [ext_mOTU_v3_26709]
Bifidobacterium species incertae sedis [ext_mOTU_v3_26696]
Desulfovibrio species incertae sedis [ext_mOTU_v3_26656]
Clostridiales species incertae sedis [ext_mOTU_v3_26617]
Clostridium species incertae sedis [ext_mOTU_v3_26695]
Alistipes species incertae sedis [ext_mOTU_v3_33183]
Sutterella species incertae sedis [ext_mOTU_v3_26637]
Clostridiales species incertae sedis [ext_mOTU_v3_26595]
Faecalibacterium species incertae sedis [ext_mOTU_v3_26928]
Bifidobacterium species incertae sedis [ext_mOTU_v3_26736]
Oscillibacter species incertae sedis [ext_mOTU_v3_26937]
Clostridiales species incertae sedis [ext_mOTU_v3_33798]
Clostridiales species incertae sedis [ext_mOTU_v3_26949]
Clostridiales species incertae sedis [ext_mOTU_v3_26523]
Clostridiales species incertae sedis [ext_mOTU_v3_26752]
Lachnospiraceae species incertae sedis [ext_mOTU_v3_26654]
1E−06 1E−05 1E−04 1E−03 1E−02 1E−01 1E+00
Abundance (log10−scale)
0 1
−log10(adj. p value)
−1 −0.5 0 0.5 1
Generalized fold change
0 0.25 0.5 0.75 1
AU−ROC
B LIFELINES
Differentially abundant features showing top 50 features
Significance
Fold change
Prevalence shift
1
$6
Oscillibacter sp.[meta_13092]
Clostridiales sp.[meta_13165]
Clostridium sp. [ref_03676]
Ruminococcaceae sp.[ext_16260]
Flavonifractor plautii [ref_02971]
Bifidobacterium longum [ref_01099]
Oscillibacter sp.[ext_17930]
Flavonifractor plautii [ref_05238]
Bacteroides dorei/vulgatus [ref_02367]
Parabacteroides distasonis [ref_03640]
Bacteroides sp. [ref_03475]
Clostridium phoceensis [ref_08865]
Oscillibacter sp.[ext_28927]
Alistipes sp.[ext_18141]
Bacteria sp.[ext_17417]
Ruminococcus sp.[meta_12711]
Phascolarctobacterium sp.[ext_27004]
Ruminococcaceae sp.[meta_12291]
Mycoplasma sp. CAG:472 [ref_07685]
Ruminococcus sp.[meta_12535]
Bacteroides sp.[ext_17856]
Firmicutes sp.[meta_13184]
Anaerotruncus sp. CAG:390 [meta_14000]
Clostridium sp.[meta_12977]
Clostridiales sp.[meta_12386]
Firmicutes bacterium CAG:238 [ref_07735]
Firmicutes sp.[ext_17797]
Firmicutes sp.[meta_12438]
Clostridiales sp.[meta_14315]
Clostridia sp.[meta_14185]
Bacteroides coprophilus [ref_04094]
Oscillibacter sp. PC13 [ref_10185]
Clostridiales sp.[meta_13308]
Firmicutes sp.[meta_12365]
Ruminococcaceae sp.[meta_12664]
Bacteroides coprocola [ref_11279]
Firmicutes bacterium CAG:313 [ref_07737]
Clostridiales sp.[ext_26655]
Clostridiales sp.[meta_13159]
Clostridiales sp.[meta_12447]
Clostridiales sp.[ext_17847]
Bacteroides sp.[ext_17779]
Ruminococcaceae sp.[ext_17414]
Clostridiales sp.[meta_12763]
Butyrivibrio crossotus [ref_03686]
Bacteria sp.[ext_16372]
Clostridiales sp.[meta_13176]
Clostridiales sp. [ref_03605]
Oscillibacter sp.[ext_17503]
Ruminococcus sp. CAG:177 [meta_12400]
1E−06 1E−05 1E−04 1E−03 1E−02 1E−01 1E+00
Abundance (log10−scale)
0 1 2
−log10(adj. p value)
−1 −0.5 0 0.5 1
Generalized fold change
0 25 50 75 100
Prevalence [%]

**C** UK Twin

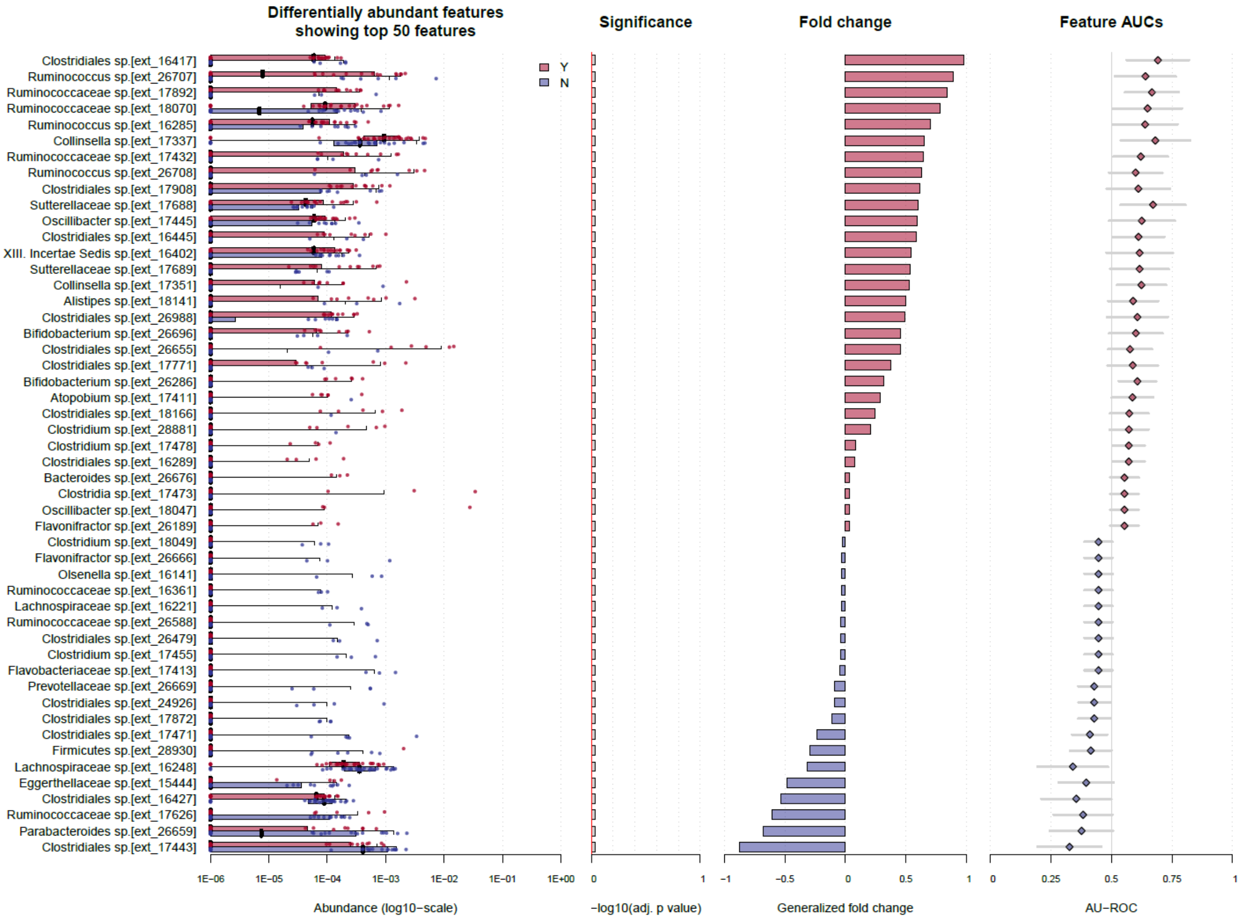


**Supplementary Figure 3.** Assessment of enriched taxa by mOTU in the gut microbiota of OA participants representing 50 differentially abundant taxa enriched in OA (red) and control (blue) in A) Estonia Biobank B) Lifelines C) UK Twin cohort

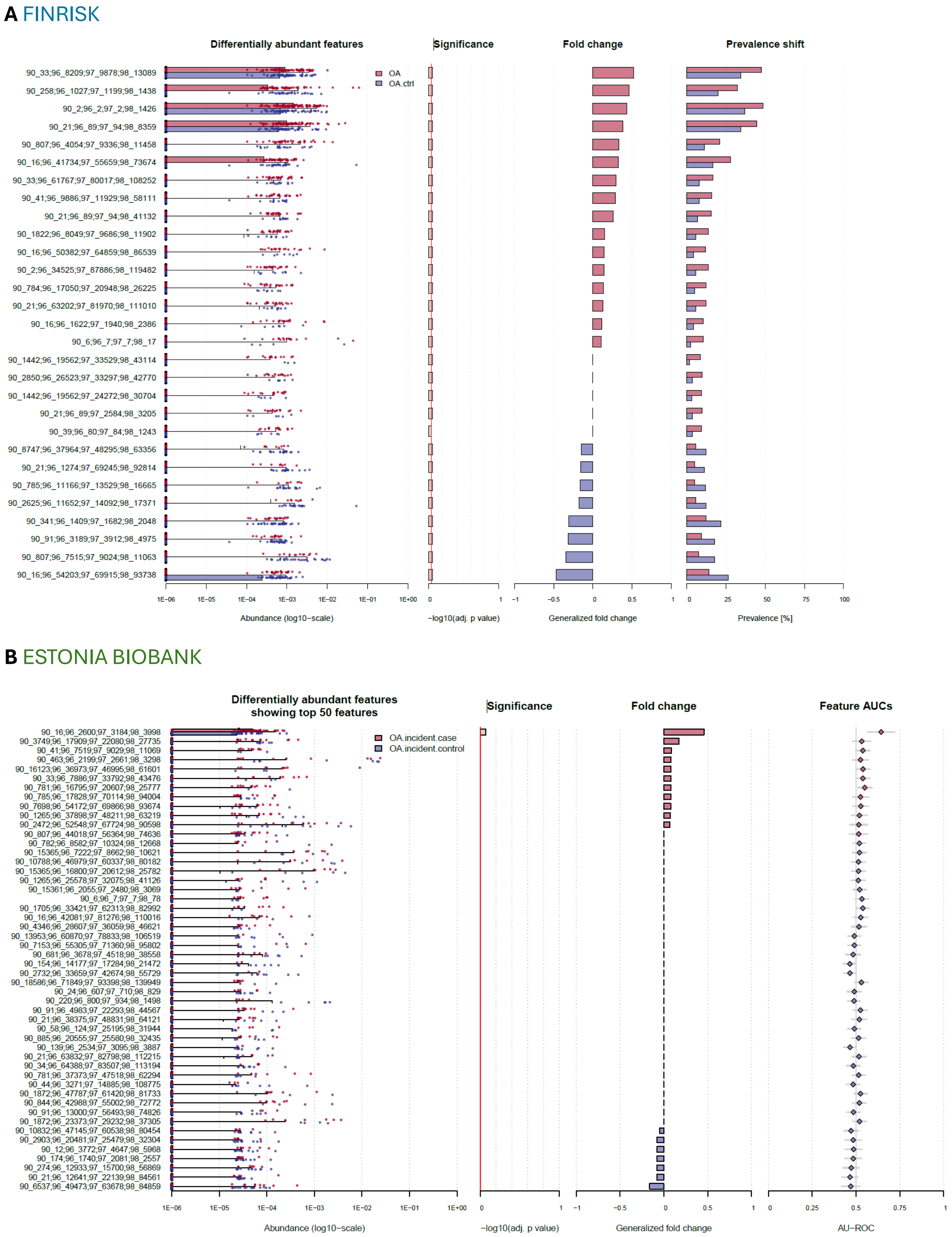
A FINRISK
Differentially abundant features
Significance
Fold change
Prevalence shift
OA
OA.ctrl
90_33;96_8209;97_9878;98_13089
90_258;96_1027;97_1199;98_1438
90_2;96_2;97_2;98_1426
90_21;96_89;97_94;98_8359
90_807;96_4054;97_9336;98_11458
90_16;96_41734;97_55659;98_73674
90_33;96_61767;97_80017;98_108252
90_41;96_9886;97_11929;98_58111
90_21;96_89;97_94;98_41132
90_1822;96_8049;97_9686;98_11902
90_16;96_50382;97_64859;98_86539
90_2;96_34525;97_87886;98_119482
90_784;96_17050;97_20948;98_26225
90_21;96_63202;97_81970;98_111010
90_16;96_1622;97_1940;98_2386
90_6;96_7;97_7;98_17
90_1442;96_19562;97_33529;98_43114
90_2850;96_26523;97_33297;98_42770
90_1442;96_19562;97_24272;98_30704
90_21;96_89;97_2584;98_3205
90_39;96_80;97_84;98_1243
90_8747;96_37964;97_48295;98_63356
90_21;96_1274;97_69245;98_92814
90_785;96_11166;97_13529;98_16665
90_2625;96_11652;97_14092;98_17371
90_341;96_1409;97_1682;98_2048
90_91;96_3189;97_3912;98_4975
90_807;96_7515;97_9024;98_11063
90_16;96_54203;97_69915;98_93738
1E−06 1E−05 1E−04 1E−03 1E−02 1E−01 1E+00
Abundance (log10−scale)
0 1
−log10(adj. p value)
−1 −0.5 0 0.5 1
Generalized fold change
0 25 50 75 100
Prevalence [%]
B ESTONIA BIOBANK
Differentially abundant features
showing top 50 features
Significance
Fold change
Feature AUCs
OA.incident.case
OA.incident.control
1E−06 1E−05 1E−04 1E−03 1E−02 1E−01 1E+00
Abundance (log10−scale)
0 1
−log10(adj. p value)
−1 −0.5 0 0.5 1
Generalized fold change
0 0.25 0.5 0.75 1
AU−ROC

**C** LIFELINES

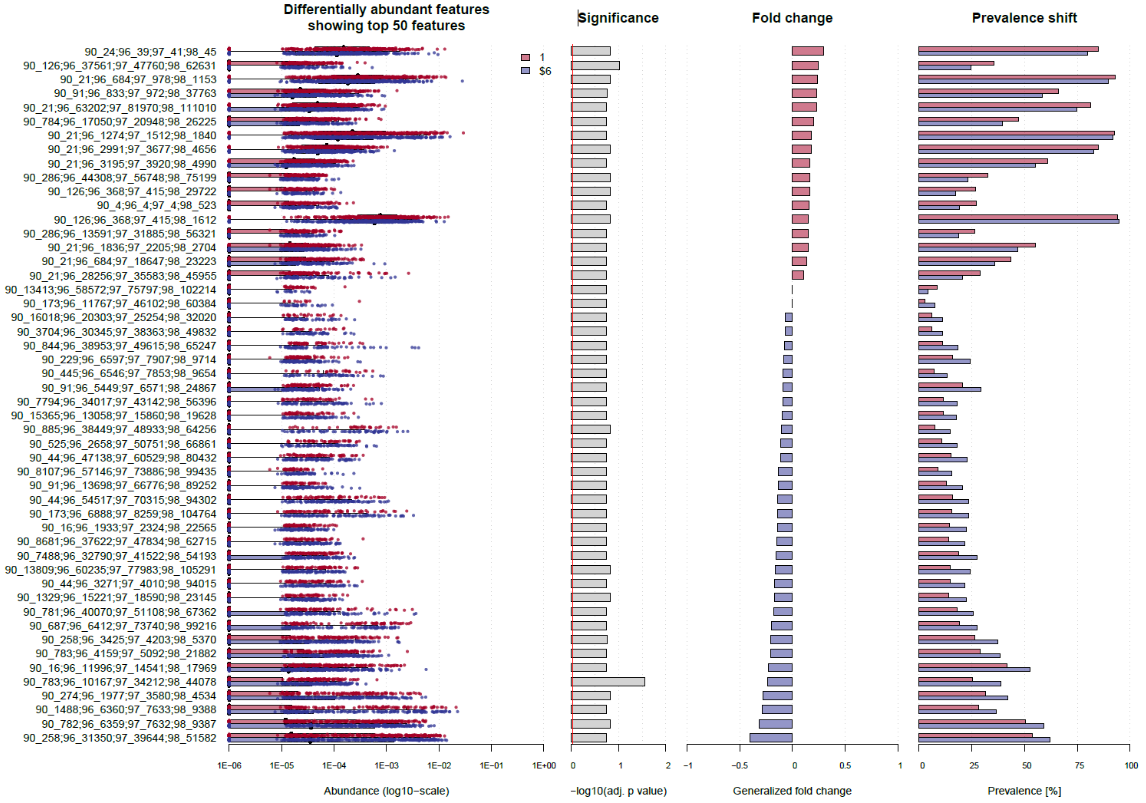


**D** UK Twin

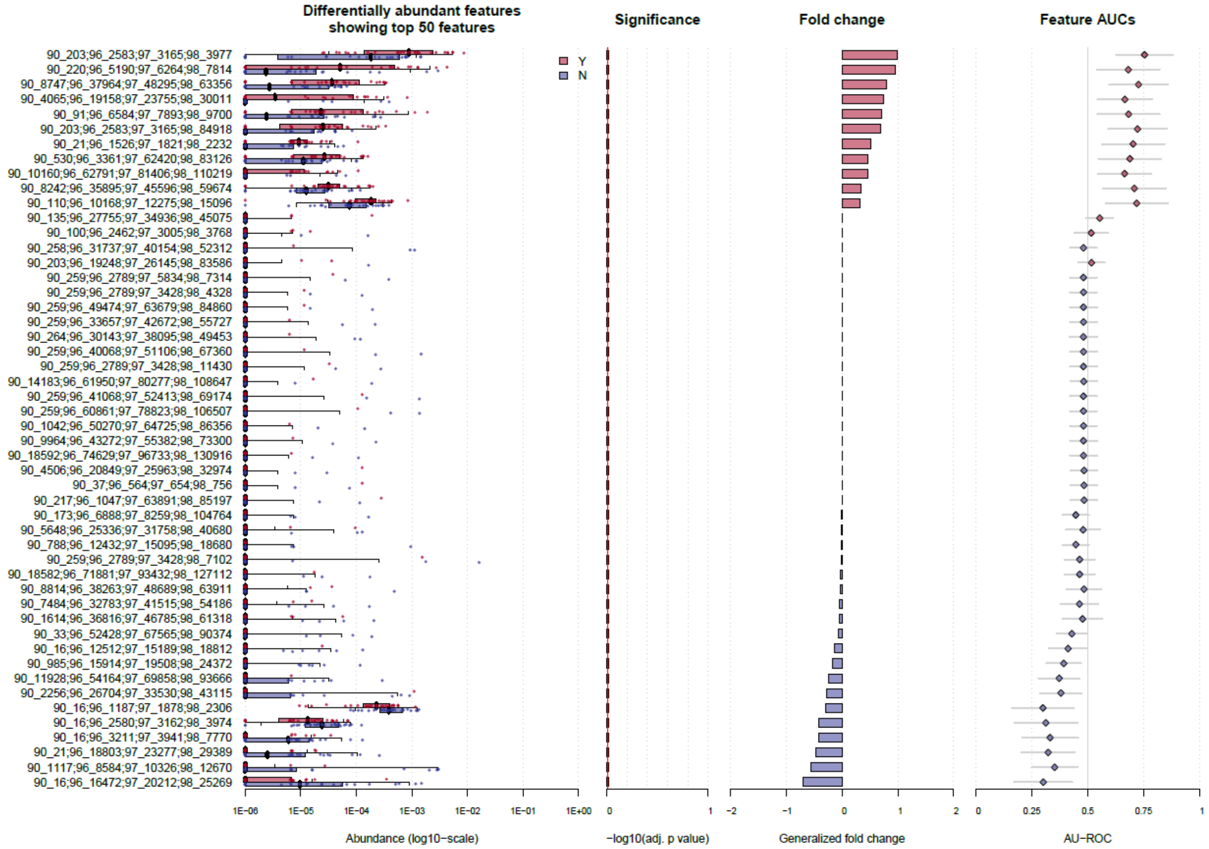

**Supplementary Figure 4** Assessment of enriched taxa by mapseq in the gut microbiota of OA participants representing 50 differentially abundant taxa enriched in OA (red) and control (blue) in A) Finrisk B) Estonia Biobank C) Lifelines D) UK Twin cohort

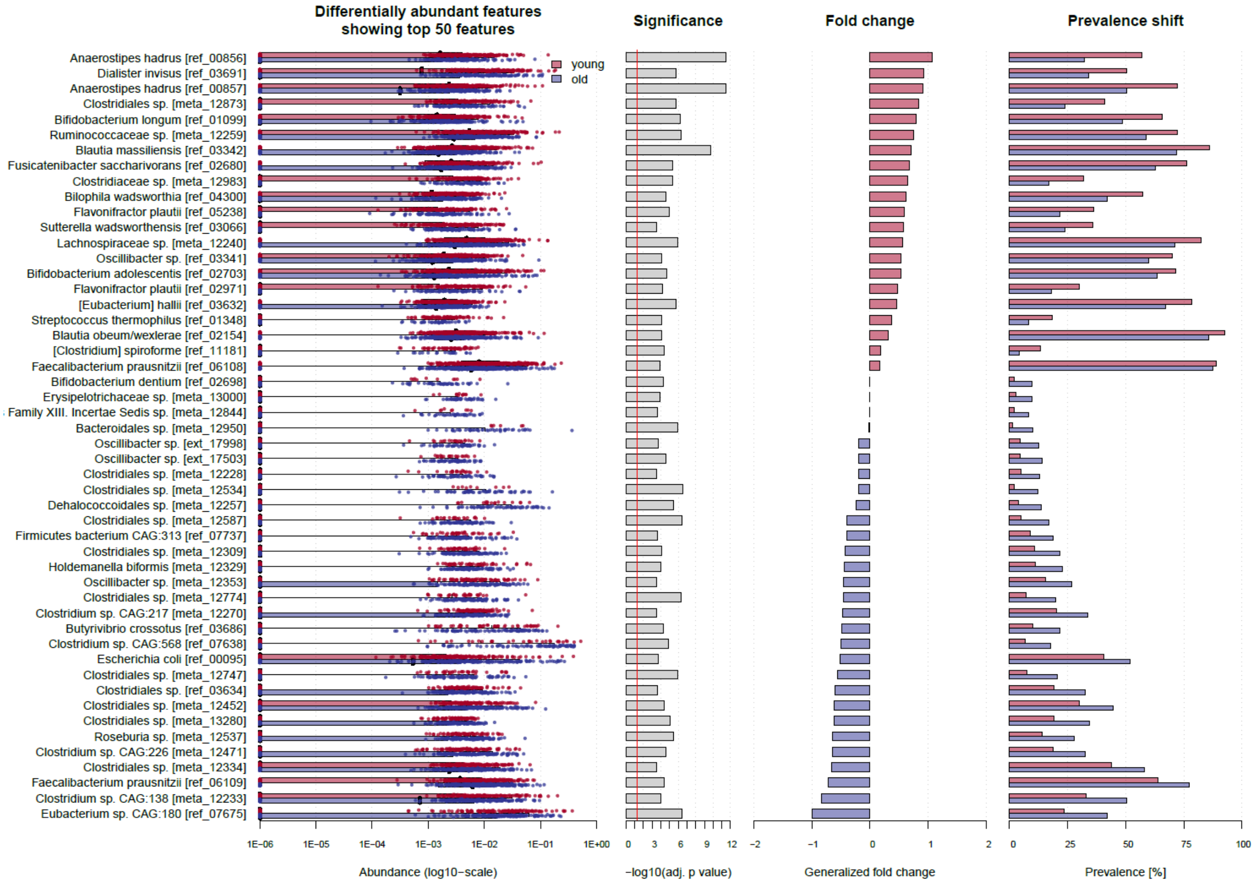


**Supplementary Figure 5** Assessment of enriched taxa by mOTU in the gut microbiota of <55 participants (red) and >65 participants (blue) representing 50 differentially abundant taxa in Finrisk cohort

**A** FINRISK HIP OA

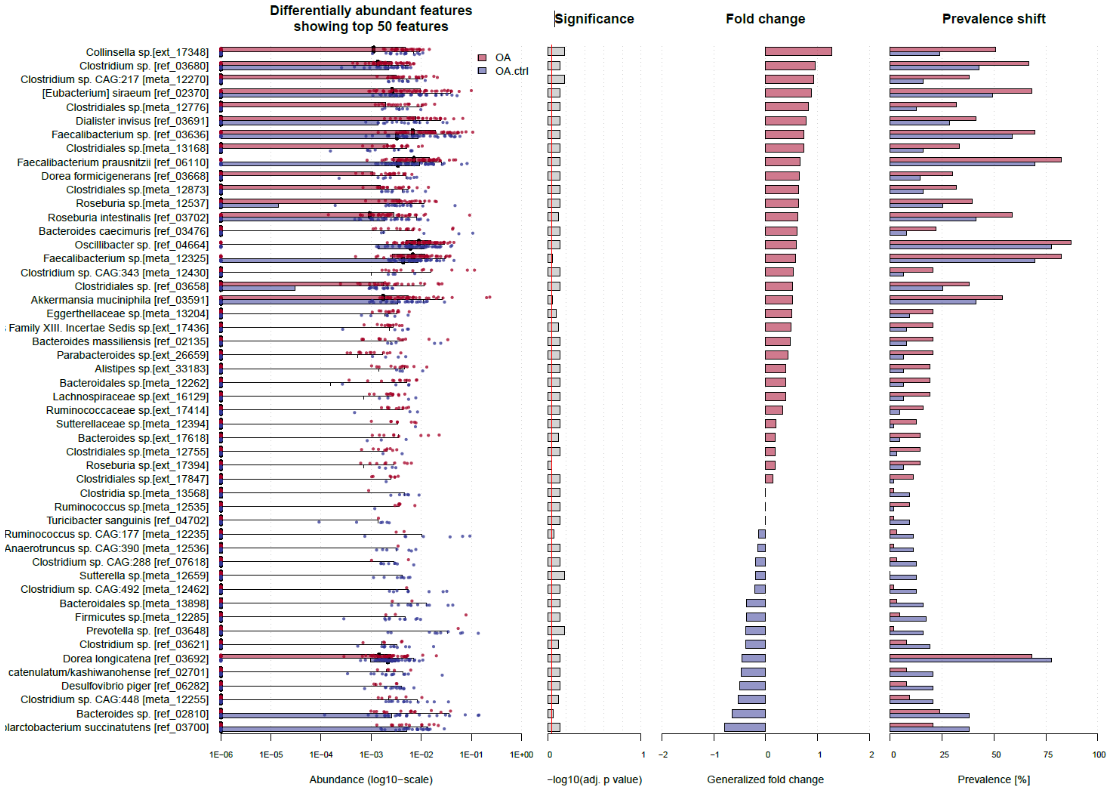


**B** FINRISK KNEE OA

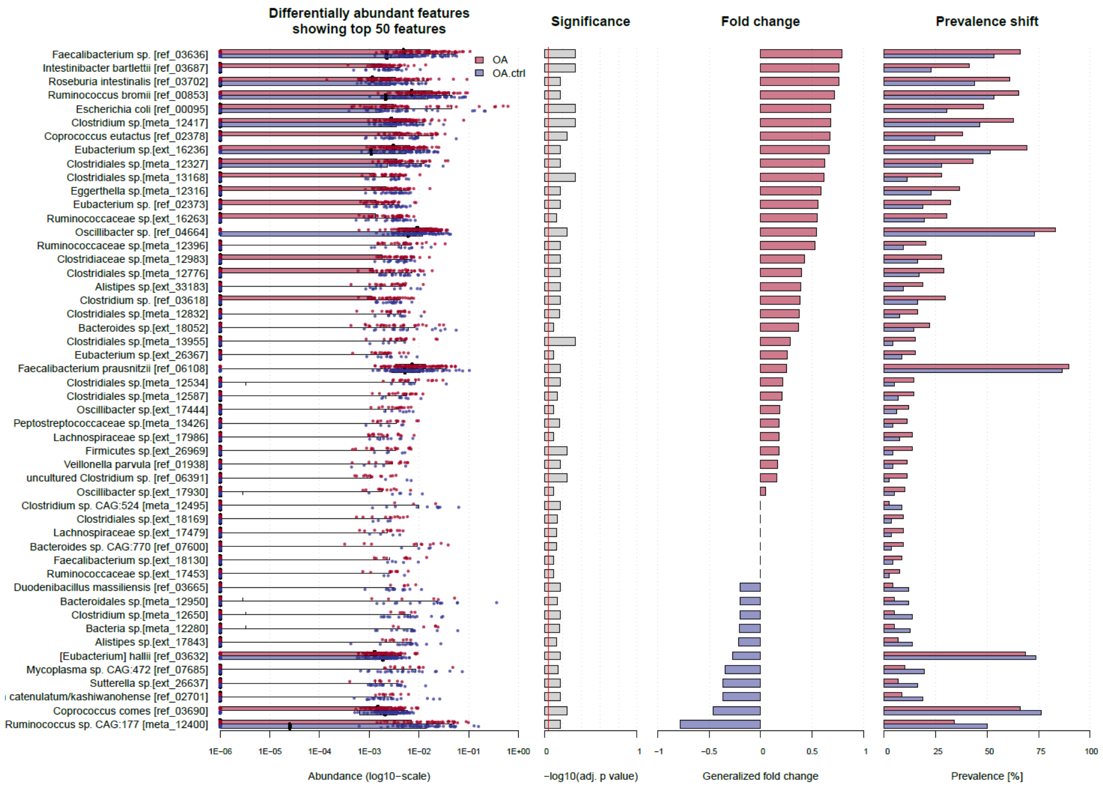

C FINRISK OTHER OA

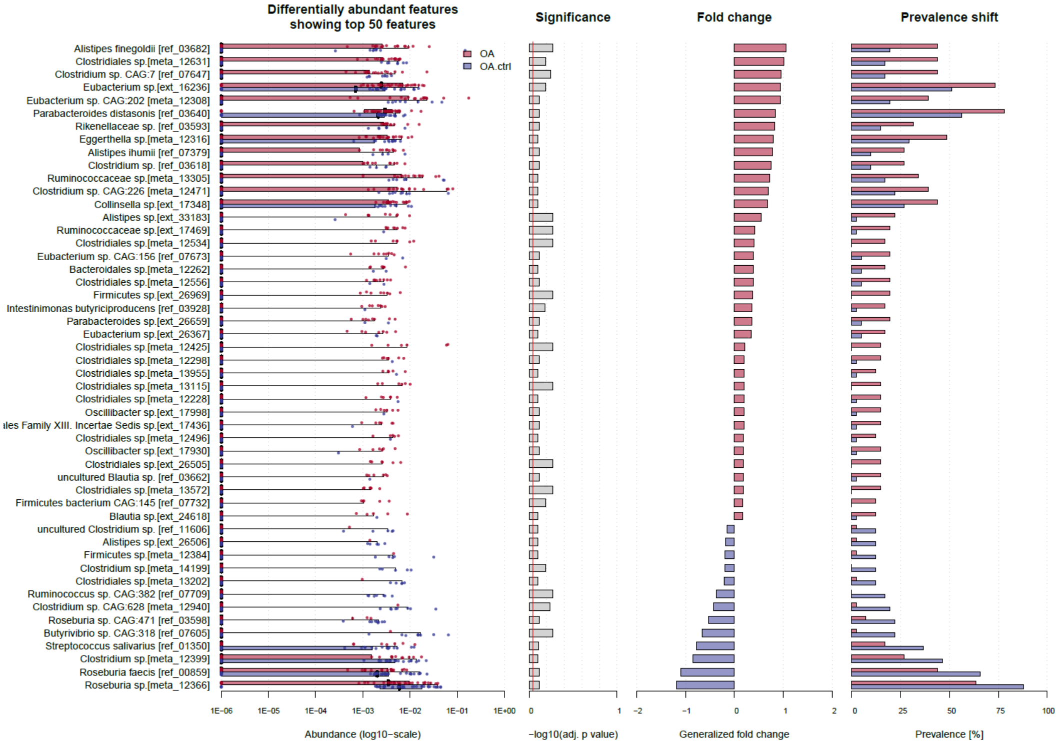


**Supplementary Figure 6.** Assessment of enriched taxa by mOTU in the gut microbiota of OA participants representing 50 differentially abundant taxa enriched in OA (red) and control (blue) in A) hip OA B) knee OA C) other OA in Finrisk biobank

**Supplementary Table 1.** The table presents the number of proteins, KEGG orthologues (KO) and Gene Ontologies (GO) present in the protein catalogue and following mapping. The column designated "proteins" enumerates the number of proteins that have been annotated for the specific annotation. The column labelled "features" lists the total number of features available in the protein catalogue. The columns labelled "UKtwins" and "EstBB" present the number of detected features after filtering as absolute and relative (in relation to 'features') numbers, respectively.

| | proteins | | features | UKtwins | | EstBB | |
|---|---|---|---|---|---|---|---|
| proteins | 12,383,493 | 100% | 12,383,493 | 1,084,913 | 9% | 1,962,482 | 16% |
| KO | 4,851,159 | 39% | 9,885 | 6,570 | 66% | 7,058 | 71% |
| GO | 521,685 | 4% | 15,406 | 8,436 | 55% | 10,424 | 68% |
